# The effects of biological knowledge graph topology on embedding-based link prediction

**DOI:** 10.1101/2024.06.10.598277

**Authors:** Michael S. Bradshaw, Anton Avramov, Alisa Gaskell, Ryan M. Layer

## Abstract

Due to the limited information available about rare diseases and their causal variants, knowledge graphs are often used to augment our understanding and make inferences about new gene-disease connections. Knowledge graph embedding methods have been successfully applied to various biomedical link prediction tasks but have yet to be adopted for rare disease variant prioritization. Here, we explore the effect of knowledge graph topology on knowledge graph embedding link prediction performance and challenge the assumption that massively aggregating knowledge graphs is beneficial in deciphering rare disease cases and improving prediction outcomes. We find that using a filtered version of the Monarch knowledge graph with only 11% of the original size results in notably improved model predictive performance. Additionally, these findings suggest that successful KG optimization depends on selecting high-quality information rather than simply maximizing the amount of data included.

## Introduction

One of the main challenges in diagnosing rare and undiagnosed diseases is the limited information about the disease’s progression and its underlying genetic causes. Many diseases are still characterized solely by the phenotype-driven method, which is hindered by symptoms often shared between clinical findings ^1^. Similarly, understanding the role of the specific genetic variant on disease mechanism is difficult when few to no other cases have ever been reported. In such cases, knowledge graphs (KGs) can be useful for applying existing biological knowledge more broadly to make inferences about poorly characterized genes and diseases ^2^. Precision medicine utilizes knowledge graphs to model interactions and relationships between diverse biological entities, such as genes, pathways, diseases, anatomy, and therapeutics. These entities are represented as nodes, while the relationships between them are represented by edges that can contain multiple types of directed or undirected connections, creating a web of interconnected biological knowledge ^3^.

Like our collective understanding of the world around us, KGs are inherently incomplete and grow as discoveries are made and published. One heavily researched approach for filling in missing information is the link prediction task, with recent emphasis on knowledge graph embedding (KGE). KGE offers several advantages over traditional methods, including edge type awareness, the use of higher-order patterns, scalability, and improved accuracy compared to feature-based approaches. These methods have been successfully applied in drug discovery tasks^4–7^. Beyond pharmaceutical applications, KGE can be directly applied to rare disease link prediction and variant prioritization. Although several research groups have successfully applied KGs and KGE link predictors in rare disease contexts, the field lacks stand-alone variant prioritization tools based on KGE link prediction ^8^. demonstrated the potential of embeddings for clustering KGs and identifying rare disease modules, while Vilela et al. ^9^ showed that KGE link prediction could prioritize variants in conditions like Autism Spectrum Disorder. To this day, neither group published a stand-alone tool for broader use.

However, KGE link predictions face several important limitations. A major technical challenge for rare disease variant prioritization is that KGE models tend to preferentially score high-degree nodes over low-degree nodes during link prediction ^4^. This bias toward well-characterized, high-degree entities is particularly problematic given that the primary objective of link prediction is to generate insights about poorly studied genes and diseases ^10^. Beyond their impact on individual analysis, degree-biased models are part of a troubling cyclical pattern in which the overemphasis of certain genes in experimental studies gives rise to topologically imbalanced KGs ^11^. Training models on these biased KGs produces degree-biased models that inflate the importance to well-studied biological entities while underestimating others ^4^. We term this phenomenon the bias cycle (**Fig. 1A**).

**Figure 1.**
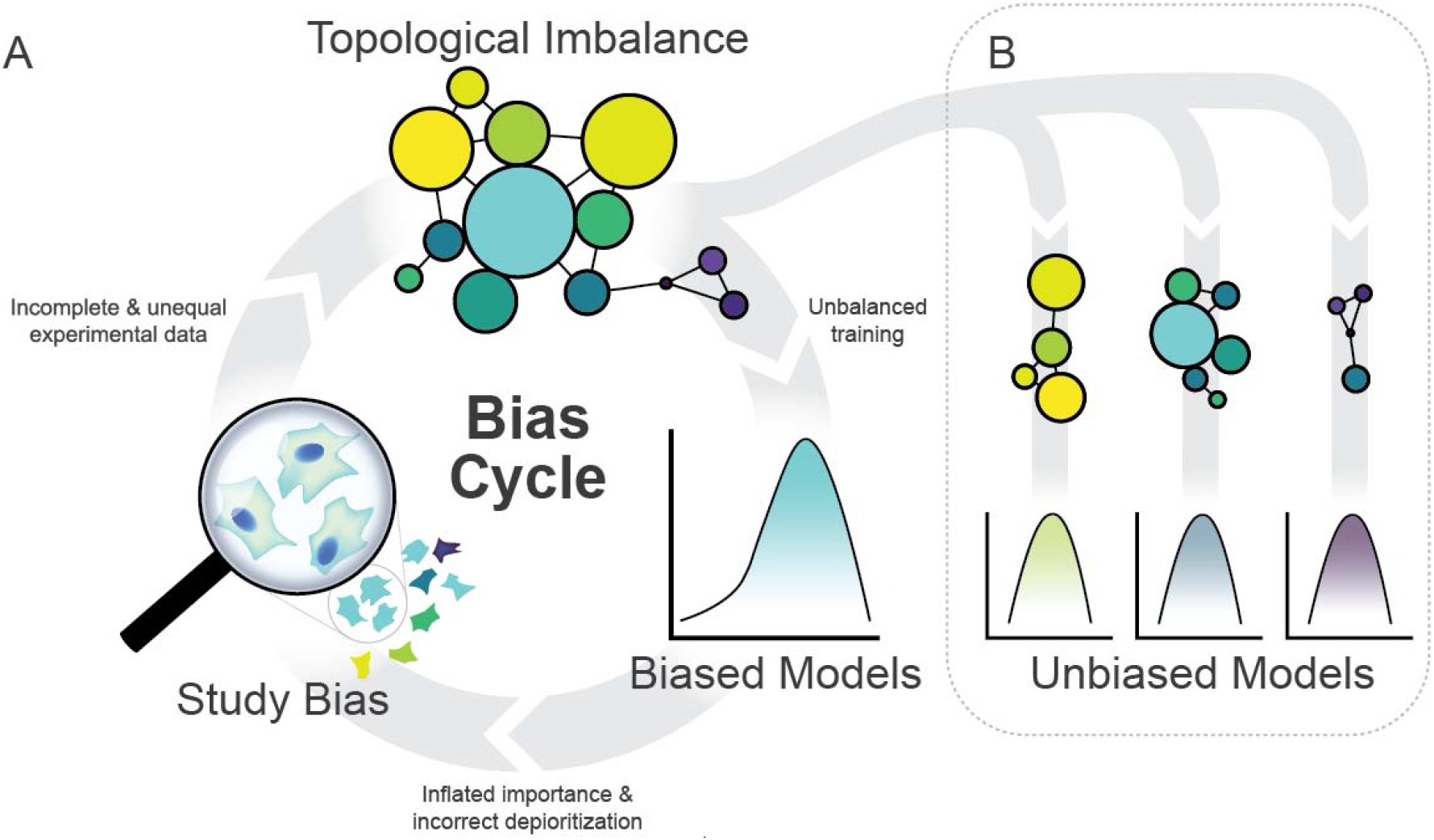
A visual description of the bias cycle. Study bias leads to incomplete and unequal data representation, which is then incorporated into KG that exhibit topological imbalance. Training predictive models on this imbalanced data produces biased models that inflate the importance of well-studied entities while systematically under ranking others. These model predictions subsequently inform future study design, perpetuating the cycle.

Importantly, topological imbalance does not originate from underlying biological nature but rather from researchers’ skewed interest in certain genes, proteins and biochemical mechanisms, which leads to the aforementioned bias ^11^. It is worth noting, however, that not all biochemical mechanisms are equally accessible to study. This creates a situation where well-studied genes and protein targets receive disproportionate attention and detailed mechanistic understanding, while others remain virtually ignored ^12–14^. Remarkably, just 19% of genes receive 49% of attention ^13^. This imbalance is particularly concerning since rare genetic disease research, although less studied, has historically provided crucial insights into human disease mechanisms ^15^.

This research bias is further reflected in the fundamental differences in data quality used for constructing KG metanodes. High quality data prioritizes semantic precision and curation, featuring carefully validated concepts with comprehensive metadata and expert oversight that ensure reliable reasoning, though at the cost of limited coverage. In contrast, high quantity data emphasizes comprehensive coverage through automated generation techniques, rapidly creating extensive concept hierarchies that capture emerging topics but often sacrifice semantic coherence due to minimal validation.

One of the key advantages of using KGs is their ability to fill gaps in our understanding of relationships between biological entities, which is essential for studying rare diseases. It is generally believed that the more information a KG contains, the better the predictive performance will be. This additional information could essentially come from any source, whether from anatomical, pharmaceutical, human, or any other model organism databases. Indeed, several high-quality KG-based methods have been developed using information aggregated from multiple data sources . However, in the context of KGE models, this approach of creating one big KG remains untested and could prove counterproductive, since topological imbalance may increase with data aggregation.

Here, we investigated the implications of degree-bias and KG aggregation when predicting connections among specific groups of genes and diseases and explored whether the assumption that more data is always better when training KGE link prediction models (**Fig 1B**). Using KGE models trained on the Monarch KG, we found that heritable cancer genes perform better than non-cancer genes in link prediction tasks, while rare diseases, ultra-rare diseases, and other disease categories lag behind heritable cancer genes in predictive performance. We show that using all types of information in a highly heterogeneous KG can degrade the performance of KGE link predictors, suggesting that the set of metanodes selected from a KG should undergo a feature selection-like process to identify what types of information are helpful for a given problem. Additionally, we found that a balance must be struck between the number and quality of edges used for training; using all available edges is suboptimal, but overly strict filtering criteria can be even worse. These findings will help improve KGE-based link predictors as the field adopts them for rare disease variant prioritization.

## Methods

To investigate KGE link prediction performance across different gene and disease categories, we optimized and trained a TransE ^16^, RotatE ^17^ and ComplEx ^18^ models on the Monarch KG ^19^. We evaluated performance across three models on link prediction tasks stratified by five groups: heritable cancer genes, non-cancer genes, rare diseases, ultra-rare diseases, and control diseases.

To test whether more data consistently improves performance, we systematically created filtered versions of the KG (**Fig. 2**). Starting with a minimal graph containing only gene and disease nodes and edges, we incrementally reintroduced one additional metanode type (from 22 possible types) (**Table S1)** and measured the impact on link prediction accuracy. Additionally, we explored the trade-off between edge quality and quantity by comparing protein-protein interaction data from HuRI (high quality) ^20^ and STRING (high quantity) ^21^ databases.

**Figure 2.**
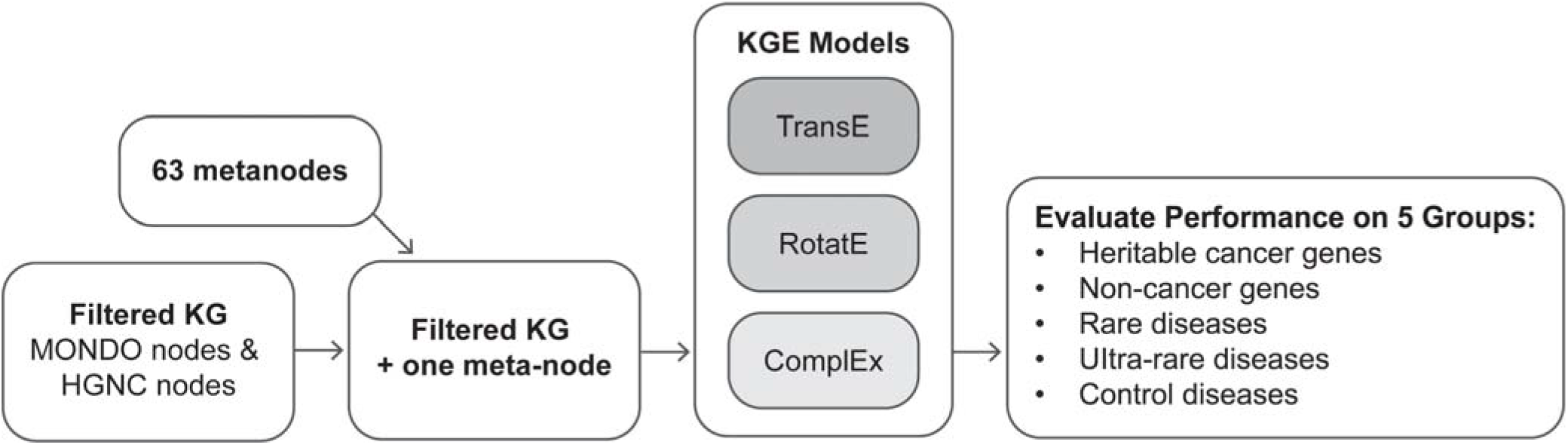
Knowledge graph embedding evaluation workflow. The pipeline begins with 63 metanodes that are filtered to create a knowledge graph containing MONDO and HGNC nodes, with an additional meta-node. Three knowledge graph embedding models (TransE, RotatE, and ComplEx) are trained on this filtered graph and evaluated for link prediction performance across five disease categories: heritable cancer genes, non-cancer genes, rare diseases, ultra-rare diseases, and control diseases.

### Knowledge graphs

### Monarch knowledge graph

For this study we used the 2023-12-16 version of Monarch KG, a large and highly heterogeneous graph containing medically relevant data about genes, proteins, disease, phenotypes, and anatomy in numerous model organisms such as human, mouse, zebrafish and fruit fly ^22^. Edges in Monarch KG are structured as semantic triples in the format subject → predicate → object, where subject and object are nodes and the predicate describes the type of edge, or relationship, connecting the two nodes. For example, NBEA → interacts_with → NOTCH1, indicating a physical interaction between the two proteins, or NBEA → causes → epilepsy indicating a causal link between a variant in the NBEA gene and epilepsy. Monarch KG has 519,303 nodes across 63 (**Fig. 3A**), and 11,412,471 edges across 23 edge types (**Fig. 3B**).

**Figure 3.**
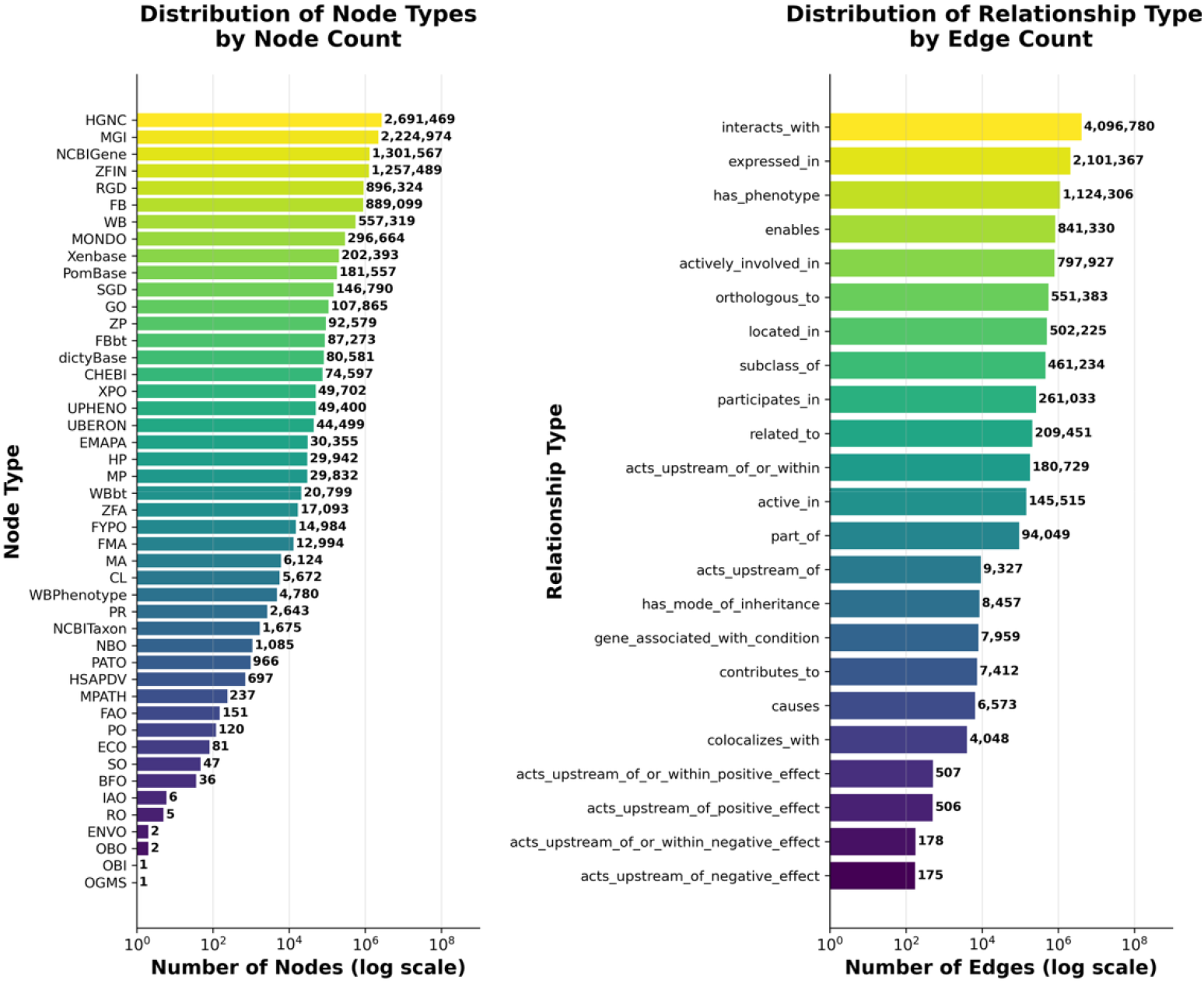
Distribution of node and edge types in Monarch Knowledge Graph. **A**. Types of nodes and their corresponding frequencies. **B**. The number of edges of each type

### Filtered knowledge graphs

To evaluate KGE models at the link prediction task, we created a set of filtered versions of the Monarch KG (hereinafter Filtered KG) by retaining only nodes and edges originating from the Monarch Disease Ontology (MONDO nodes) and HUGO Gene Nomenclature Committee identifiers (HGNC nodes) ^23^ resulting in a 92% reduction in edge number (**Fig. 2, Table S1**). We identified 22 metanodes from the full KG that were connected to the HGNC and MONDO nodes in our filtered graph and created a new filtered KG for each of these additional metanode types.

### HuRI

HuRI is a protein-protein interaction (PPI) network derived from a large-scale “all-by-all” binding experiment that should be free from the experimental study design bias that affects aggregate PPI networks ^11,20^. We hypothesized that using this unbiased PPI data to train KGE models would improve predictive performance by reducing the research bias present in literature-aggregated networks. Thus, we created a version of the filtered KG where all HGNC nodes and edges were removed and replaced with those from HuRI, which resulted in a graph with 39,120 nodes and 6,210,908 edges.

### STRING

We also constructed an additional version of the filtered KG using additional edges and nodes from a high quantity PPI network STRING ^21^ (**Table S1**). STRING is an aggregate PPI network that derives link information from 3^rd^ party databases, experimental results, co-expression networks, and text-mining. STRING also provides edge confidence scores describing the amount of evidence behind an edge, stratified by the type(s) of sources it originates from. Working with the human version of STRING 12.0, we removed all edges that are not experimentally derived and based on the confidence scores we created three additional KGs containing the top 25%, 50%, and 100% most confident edges, which replaced all HGNC nodes and edges.

### KGE models and optimization

TransE, RotatE, and ComplEx knowledge graph embedding models were optimized and trained for each KG version using the PyKEEN package ^24^. Although the KGE models used in the current study were developed over a decade ago, they remain an industry standard for the link prediction tasks and continue to be widely applied in recent research ^25–28^. For each KG, edges were randomly divided into an 80/10/10 train/validation/test splits. Hyperparameters were optimized using quasi-Monte Carlo (QMC) sampling ^29^ to select parameters from the ranges listed in Supplemental Table 2. Parameter ranges were informed by previous work on these models in a similar learning task ^30^. Hyperparameter optimization for each model was run on a single dedicated NVIDIA A100 GPU with 80GB VRAM. Trial run times were collected from the hyperparameter optimization trial logs for all 30 iterations for each model-KG combination.

### Datasets

#### Heritable cancer genes

We compiled a list of 25 genes with mutations linked to hereditary cancer risk associated with breast, colorectal, endometrial, fallopian tube, ovarian, primary peritoneal, gastric, pancreatic, skin, and prostate cancer from FORCE - Facing Our Risk of Cancer Empowered (FORCE, n.d. https://www.facingourrisk.org).

#### Non-cancer genes

We randomly selected 500 genes from all HGNC genes, excluding those in the heritable cancer gene set.

#### Rare and ultra-rare diseases

We obtained disease prevalence data from the July 2023 Orphanet Epidemiology report (https://www.orphadata.com/data/xml/en_product9_prev.xml). We applied the following filters: “PrevalenceGeographic” = “Worldwide,” “PrevalenceValidationStatus” = “Validated,” and diseases with cross-references to MONDO identifiers in our Monarch KG version. For ultra-rare diseases (prevalence < 1/1,000,000), this filtering yielded 3,007 eligible disorders, from which we randomly selected 300. For rare diseases (prevalence > 1/1,000,000), the same criteria yielded 290 diseases, all of which were included.

#### Control disease group

We randomly sampled 300 MONDO disease identifiers to serve as control group.

## Results

We trained TransE, one of the most widely used KGE models, on the full Monarch KG and observed evidence of node-degree bias. Of the five groups of genes and diseases we tested, link prediction for non-cancer genes, rare diseases, ultra-rare diseases, and control diseases performed poorly relative to heritable cancer genes, highlighting how baseline KGE model would be ill-suited for rare disease variant prioritization. RotatE and ComplEx, which are more robust in handling different types of information, improved performance but still exhibited the same pattern, with the four non-cancer groups lagging well behind heritable cancer genes. Interestingly, models trained on filtered minimal versions of the Monarch KG improved predictive performance for all gene groups and some disease groups, demonstrating that using all available information in a KG can be detrimental to performance (**Figs. 5, S1, S2**). Reintroducing certain metanodes to the filtered KG further enhanced performance, revealing the need to systematically explore the metanode combination space (**Fig, 6**). Finally, experiments examining the quality of PPI edges (**Fig. 7**) showed that there is an optimal balance between using all available edges (including lower quality ones) and using fewer high-quality edges.

### Degree bias

All KGE models we trained on unfiltered Monarch KG exhibited degree bias. The degree of a gene and its link prediction score had a Pearson correlation coefficient of 0.48 (median 0.20), 0.55 (median 0.37), 0.42 (median 0.23) for TransE, RotatE, and ComplEx, respectively (**Fig. 4A**). Notably, the degree bias we observed was less pronounced than that same model trained on Hetionet ^4^.

**Figure 4.**
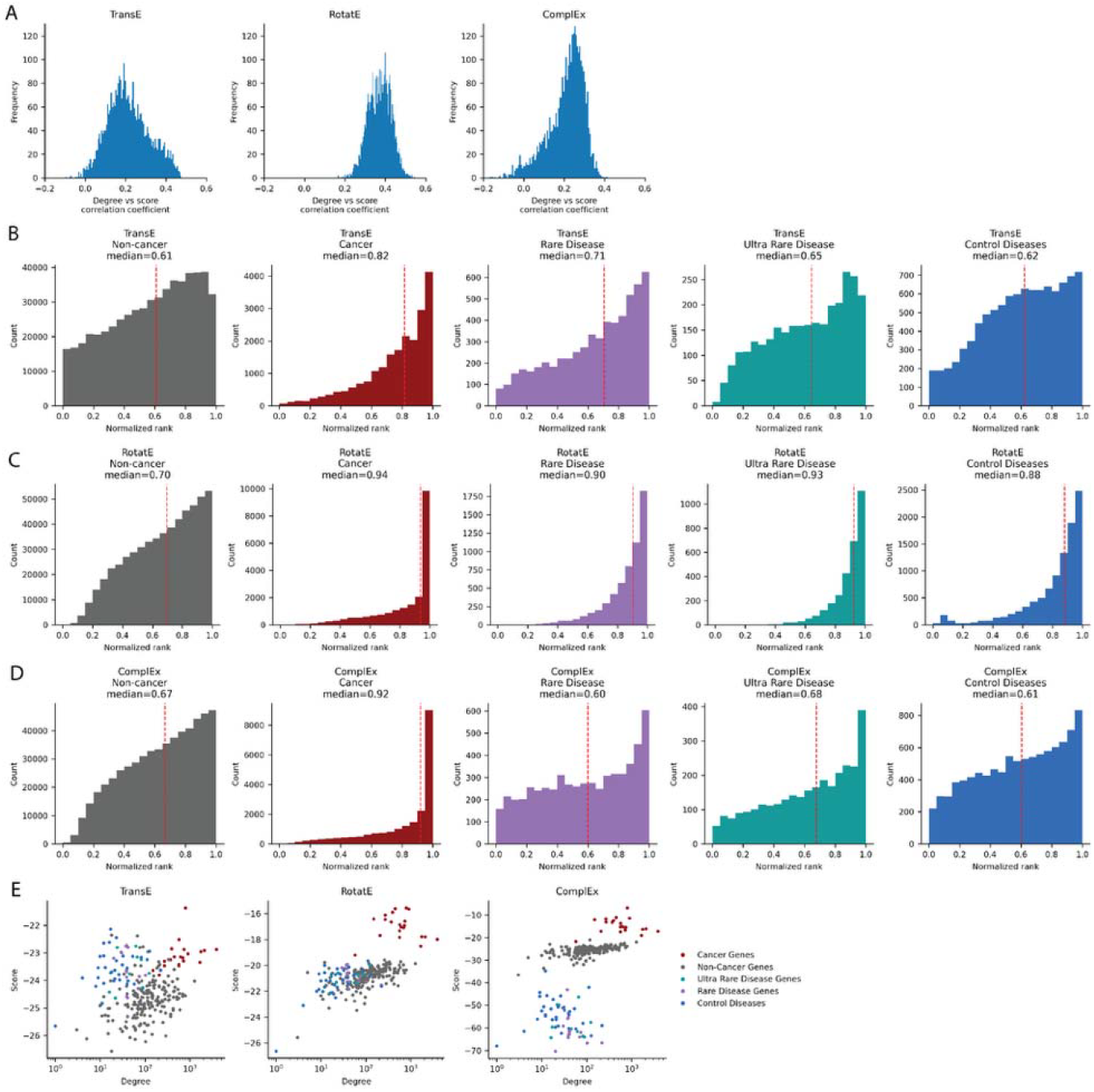
Degree bias analysis across KGE models. **A**. Distribution of correlation coefficients of the relationship between node degree and predicted association scores. Scores between genes and MONDO disease were generated in pair-wise fashion, each score representing the likelihood of a connection existing between the gene and disease – from which, correlation coefficients were calculated. **B-D**. Distribution of normalized performance scores (1-MNR) for test set predictions for **B**. TransE, **C**. RotatE, and **D**. ComplEx models. **E** Relationship between degree and score in test set edges divided by groups and models.

While TransE performed poorly across most gene and disease groups – with median normalized rank (1-MNR) of 0.61 for non-cancer genes, 0.71 for rare diseases, 0.65 for ultra-rare diseases, and 0.62 for control diseases - performance was notable better for heritable cancer genes (1-MNR = 0.82) (**Fig. 4B**). RotatE showed consistent improvements over the TransE across all groups: +12% for cancer, +19% for rare diseases, +28% for ultra-rare diseases, +19% for non-cancer genes, and +26% for control diseases (**Fig 4C** and **D**). This highlights the importance of careful model selection. Regardless of the model used, heritable cancer consistently achieved better link prediction performance than other groups genes (**Fig. 4E**).

Consistent with previous findings ^10,11^, node-degree biased models successfully ranked well-studied, higher-degree nodes but often failed to correctly prioritize results for less studied entities such as non-cancer genes, rare and ultra-rare diseases, and genetic diseases at large. This creates a problematic feedback loop: when these skewed results inform experimental design decisions, they perpetuate existing research biases and further marginalize understudied disease areas.

### Knowledge graph filtration

Using unfiltered Monarch KG for training was detrimental to predictive performance and computation run time. The full Monarch KG contains 63 metanodes representing the relationships between diverse entities, including diseases, phenotypes, genes, proteins, anatomy, chemicals/drugs, and more across both humans and model organisms. This complexity contributes to degree ias where the correlation between gene degree within gene/protein interaction networks and gene degree across the entire KG (Pearson’s R = 0.34) suggests that already highly connected nodes become even more connected in the aggregate network, potentially exacerbating the effects of degree bias. To investigate if this aggregation was affecting predictive performance, we created a minimal filtered KG containing only essential metanodes required for link prediction task - HGNC gene nodes and MONDO disease nodes.

Retraining all three KGE models using minimal filtered KG resulted in substantial improvements for link prediction performance compared the original TransE model trained on full KG version. For cancer predictions, the filtered KG improved performance by 9-10% across all models, while non-cancer genes improving by 6-14% (**Fig. 5A-C**). However, disease group predictions variable results with predictions for ultra-rare disease improved with the TransE and RotatE model by 6-12%, while rare diseases and control diseases generally performed better with the full KG, except for a marginal 1% improvement in control disease group using RotatE with filtered KG.

**Figure 5.**
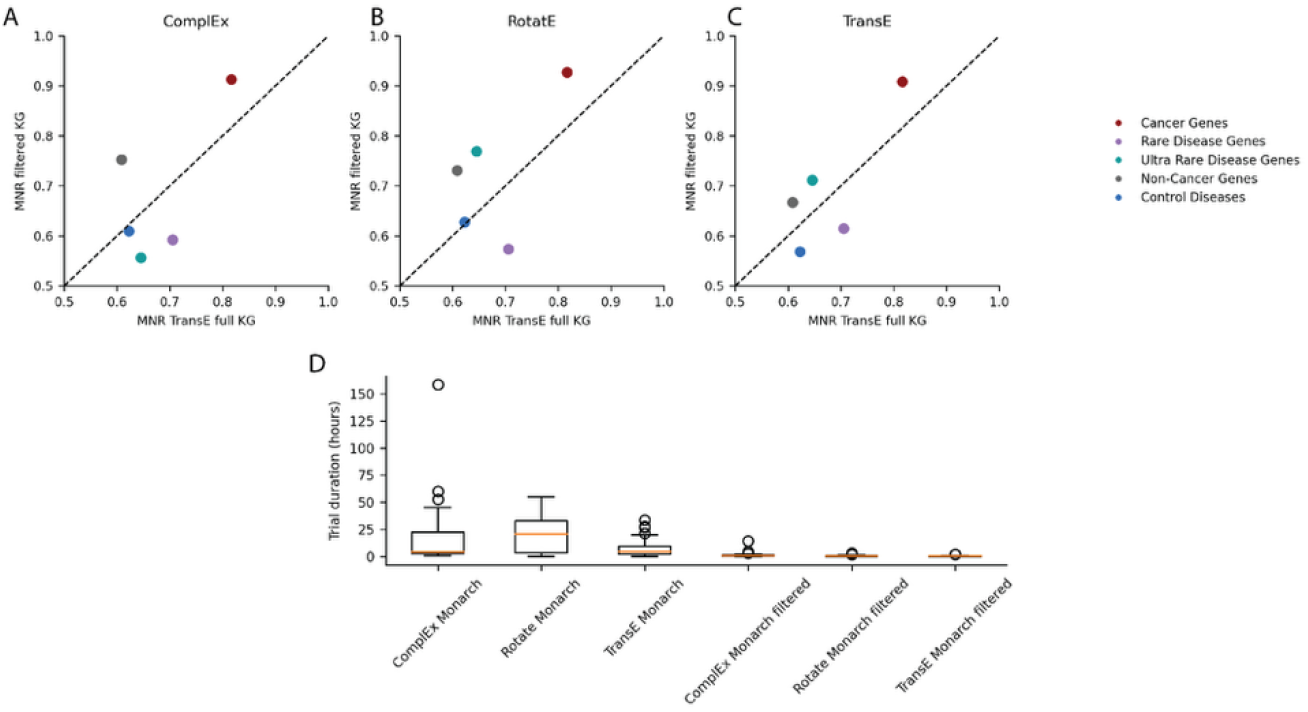
Performance and efficiency gains from KG filtering. **A-C** Comparison of performance scores (1-MNR) between filtered KG models and TransE full KG model for **A**. TransE, **B**. RotatE, and **C**. ComplEx models. X-axis shows TransE model performance trained on full Monarch KG; y-axis shows performance with filtered KG. Points above the diagonal line indicate better performance with filtered KG; points below indicate better performance with full KG. **D**. Training time per hyperparameter optimization trial, comparing full KG versus filtered KG models.

Another benefit of filtering the KG was the dramatic reduction of the computational time required for model optimization and training (**Fig. 5D**). Filtered KG models needed only 0.8%-13.8 of the training time compared to their full KG counterparts, taking hyperparameter optimization down from over 30 days to less than a day.

### Metanode reintroduction to knowledge graph

Reintroducing the Human Phenotype Ontology (HPO) to the filtered minimal KG increased performance on rare, ultra-rare, and control diseases by 10-22%. These results, together with the benefits observed from the initial KG reduction (from 63 to 2 metanodes), indicated that neither the full KG nor the minimal KG represents the optimal configuration. This suggests that an optimal combination of metanodes exists for link prediction tasks. As a first step toward identifying this optimal combination, we systematically reintroduced individual metanodes to our filtered KG.

Among the 63 metanodes in the full Monarch KG 22 are connected to the HGNC and MONDO nodes that comprise our filtered KG. Systematically reintroducing selected metanodes from this subset improved predictive performance (**Fig. 6A**). However, the magnitude of improvement varied substantially across metanodes. Reintroduction of Saccharomyces Genome Database (SGD) ^31^ provided only marginal gains, with heritable cancer genes showing the largest improvement at 3%. Similarly, dictyBase integration ^32^ provided only 1% improvement for non-cancer genes. In contrast, all disease groups (rare, ultra-rare, and control) gained significant performance improvement (10-22%) when HPO was reintroduced. Importantly, the performance gain did not correlate with the number of edges added to the network (**Fig. 6B**). For example, HPO reintroduction increased MNR by 23% in rare diseases while adding 110% more edges, whereas Chemical Entities of Biological Interest (CHEBI) ^33^ provided similar improvements with only 80% more edges. Oppositely, NCBIGene added 149% more edges but decreased performance for non-cancer genes, ultra-rare, and control diseaes. The finding that the reintroduction of HPO significantly increased performance for all disease groups is not surprising, given that it was specifically designed to standardize phenotypic descriptions of human diseases, highlighting its continued importance for computational disease analysis.

**Figure 6.**
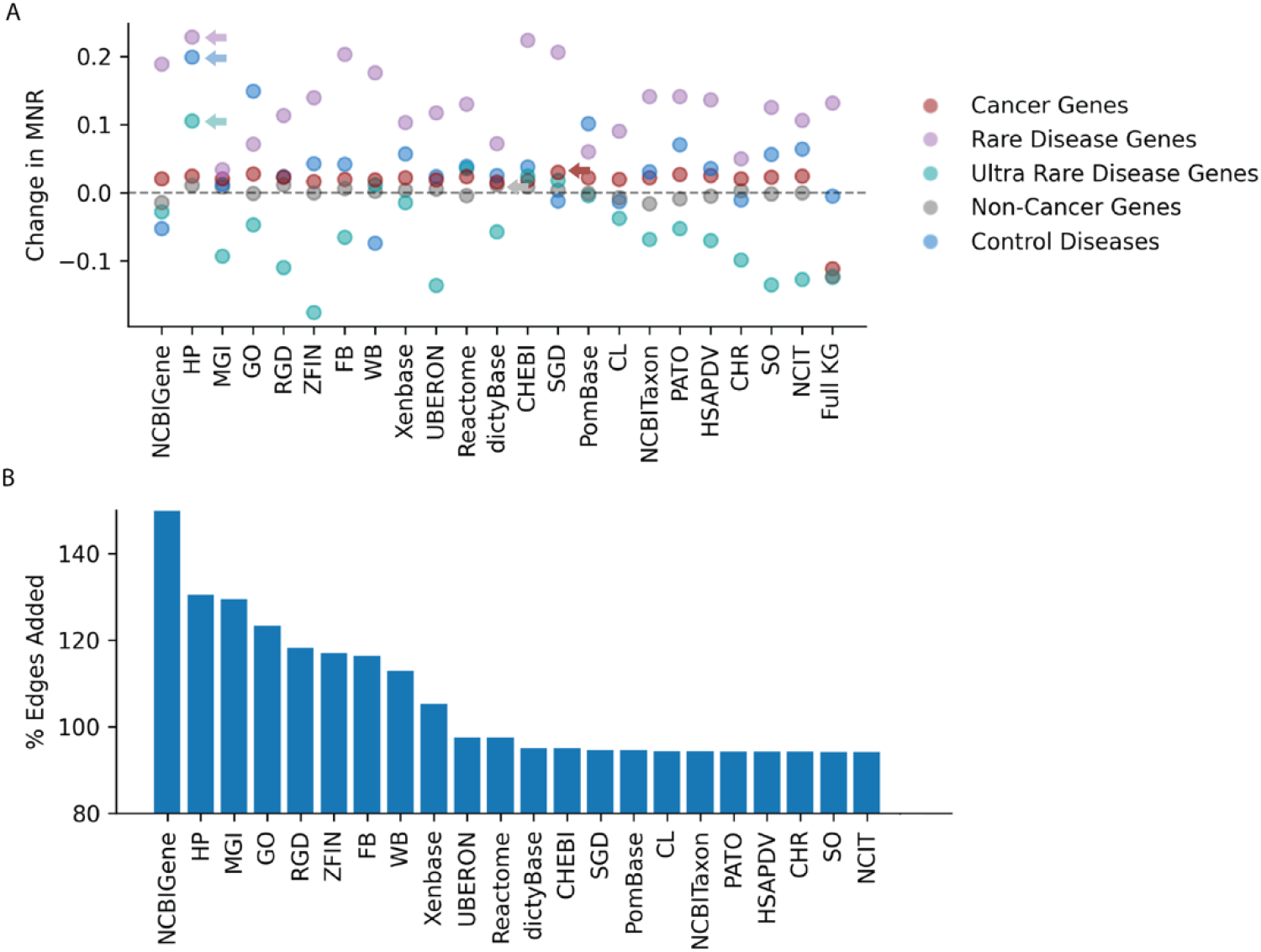
Impact of metanode reintroduction on model performance and network complexity. **A** Change in MNR when each of the 22 metanode was added back into the KG and trained on. Changes are relative to the RotatE + filtered KG model, only RotatE models were trained and reported. **B** The percent of edges added into the network by each metanode. The number of edges added by including the metanode divided by the filtered Monarch KG edge count (1,603,267).

These results suggest that optimal KG composition depends on the specific prediction task and target entities. For ultra-rare diseases, only 5 of the 22 tested metanodes improved performance when reintroduced, highlighting the need for careful metanode selection rather than bulk data aggregation.

### PPI edge filtration

Recent findings by Blumenthal et al. ^11^ challenge the reliability of PPI, showing how aggregating many small, biased studies distorts network topology. To test the hypothesis that all-by-all PPI network could control for study bias and improve prediction performance, we replaced all the HGNC nodes and edges in Monarch KG with those originating from the HuRI network.

Interestingly, use of HuRI network resulted in overall detrimental effect on performance. For non-cancer genes, MNR decreased by 16% and 17% in TransE and ComplEx respectively, while RotatE showed 5% increase. Cancer genes exhibited smaller decrease of 3% and 8% for TransE and ComplEx, with a 2% increase for RotatE. Rare diseases followed a similar pattern, 4% decrease in TransE, 6% in RotatE, but 7% improvement for ComplEx. Prediction for a control disease universally improved by 8-13%. Ultra-rare diseases could not be evaluated due to excessive node removal.

This variability and loss in performance likely originates from the drastic in edge count. HuRI contains only 3% of the edges in the full Monarch KG and includes only 10,000 proteins as opposed to the 21,000 in Monarch KG.

To test whether the performance resulted from excessive edge removal, we created a series of KGs with varying degrees of edge filtration based on experimentally derived edge confidence scores from the STRING database. We systematically reintroduced top 25%, 50%, and 100% of highest quality edges, which correspond to 15%, 36% and 71% of all PPI edges in Monarch KG. We found that peak MNR performance for non-cancer gene prediction was achieved with only 36% of the original edges (corresponding to the top 50% of STRING edges) in both ComplEx and RotatE models, while TransE MNR continually decreased as the percent of edges shrunk, suggesting that reducing the KG to only 3% of edges with HuRI was overly restrictive (**Fig. 7**). We also found that performance could be improved by removing less confident edges from the system.

**Figure 7:**
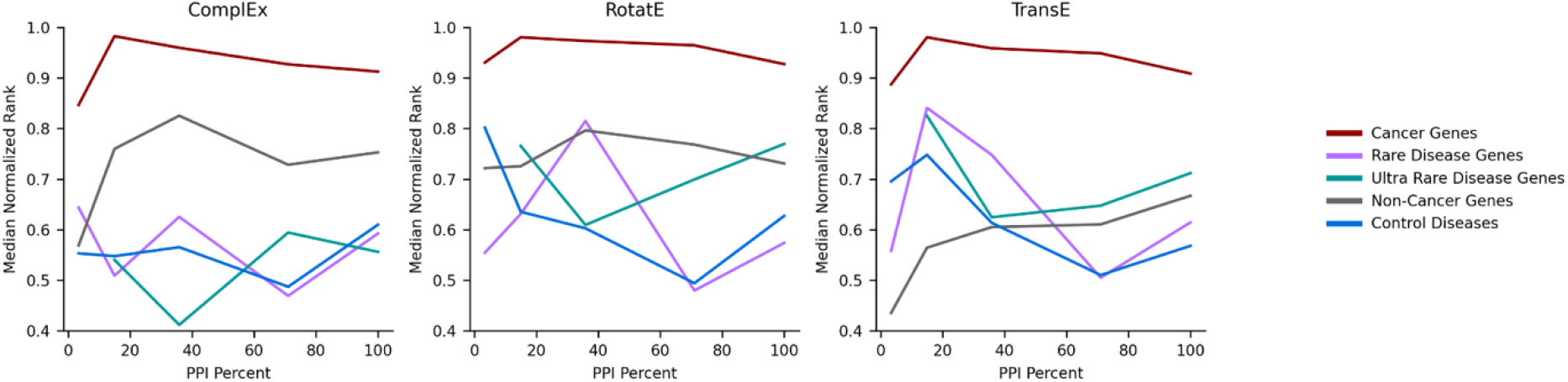
Median normalized rank performance across different gene and disease groups by embedding model. **A**. Complex, **B**. RotatE, **C**. TransE. The x-axis represents the proportion of HGNC-to-HGNC edges retained in the training set to the full Monarch KG training split.

### Performance evaluation at clinically relevant prediction thresholds

Among all the combinations of metanodes and models we evaluated, the two best-performing combinations for predicting rare and ultra-rare diseases were RotatE + Full Monarch KG (MNR = 90 and 93) and RotatE + Filterer KG with HPO re-added (MNR = 80 and 88). While MNR provides a high-level view of model performance across all edges, realistically only edges ranked in the top 100 would ever be considered for further research and evaluation, which corresponds to normalized rank > 0.99. When examining only the top-ranked k edges (up to k = 100), we found that RotatE trained on the filtered KG with HPO re-added substantially outperformed the full KG model at predicting rare disease edges (7.7% of edges were rediscovered a k=100 compared to 5.6%) despite having a lower overall MNR (**Fig. 8**).

**Figure 8:**
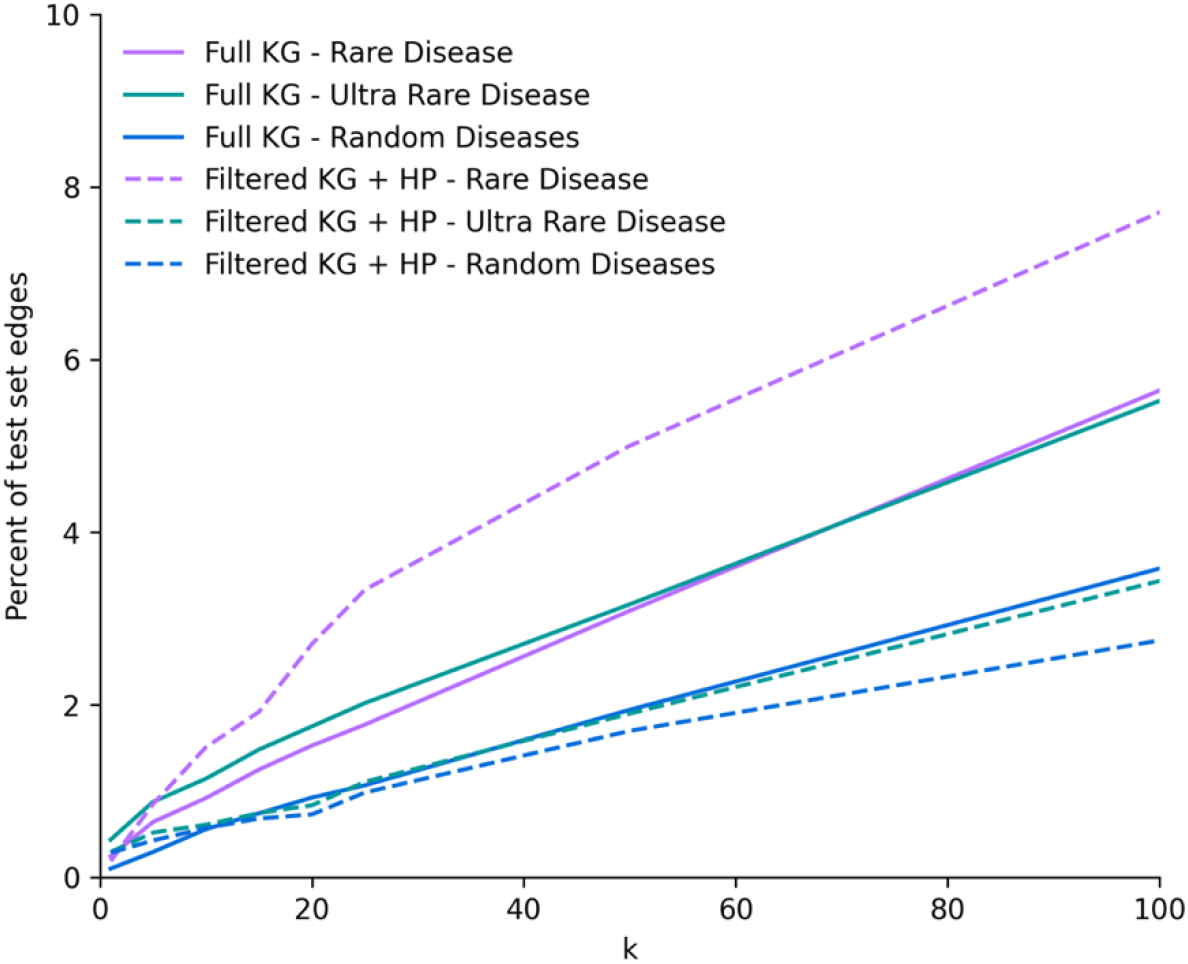
Percent of test edges ranked in the top k predictions. Dashed lines are the results for Rotate trained on the filtered KG with HPO re-added; solid lines are from RotatE trained on the full KG.

## Discussion

The disproportionate scientific attention given to a small set of genes ^13^ creates topological imbalances in knowledge graphs ^11^ that subsequently produce degree-biased predictive models ^4^. In this study we demonstrated that degree-biased models struggle to prioritize understudied entities like rare diseases correctly; however, the predictive performance and runtime can be improved through filtering the information used for training. Without such improvements, relying on degree-biased KGE models risks prioritizing already well-characterized genes and perpetuating the bias cycle.

Our results demonstrate that the quality and type of information included in a KG matter more for KGE link prediction performance than the sheer volume of data. In our metanode re-addition experiments, the most substantial performance gains came from incorporating highly curated information sources, particularly HPO.

High-degree nodes from over-studied genes and diseases create a misleading impression of strong model performance in link prediction tasks ^10^. This overall performance metric masks significant disparities across different node types, making it essential to evaluate sparse and well-connected regions of the KG independently. Models that struggle with predictions in sparse regions have limited practical value, particularly for rare disease research where reliable predictions are most needed. The real challenge lies not in refining knowledge about well-characterized genes with existing therapeutics, but in generating trustworthy predictions about poorly understood genes and diseases that currently lack approved treatments.

Despite exploring only a small fraction of all possible combinations (2.3 ^*^ 10^18^) among the 63 metanodes in Monarch KG we found that not all the information in the KG is beneficial in training and prediction performance. Our results demonstrate that superior performance and computational efficiency can be achieved with a carefully selected subset of original metanodes. While it might seem intuitive that the graph-based learning models would learn to ignore irrelevant graph structures, this assumption proves incorrect ^34^. Therefore, developing systematic methods for exploring metanode combination space to optimize KG filtering represents a promising and underexplored research direction. Ratajczak et al. ^35^ explored KG filtration by removing “counterproductive” nodes, achieving a 60% reduction in KG size and 40.8% performance improvement. However, this filtering increased the average node degree by 65% (from 24 to 49), indicating that the removed nodes were predominantly low-degree entities. While such approaches excel at improving predictions for well-studied genes and diseases, they are counterproductive when the goal is to predict less-studied entities. Our findings align with aforementioned results showing that models biased against low-connectivity nodes actually hinder progress in discovering new therapeutic targets rather than advancing it ^10^.

Given that understudied genes and diseases often intersect with health inequities, we additionally investigated whether KGE models exhibit disparities in predictive performance between male and female differentially expressed genes and between diseases caused by ancestry-specific variants in European, African, East Asian, and Latino populations. We found no significant performance differences between male and female differentially expressed genes, nor between diseases associated with different ancestry-specific variants (**Fig. S1, S2**). However, all these groups performed poorly compared to well-characterized genes such as heritable cancer genes. Notably, substantial annotation disparities exist across ancestry groups, with European ancestry variants having 2.5, 2.2, and 1.6 times more annotations than African, East Asian, and Latino ancestry variants, respectively.

We also tested the common assumption that KG prediction performance correlates with the amount and diversity of information used for training. Contrary to this expectation, we found that predictive performance improved when using a drastically filtered version of the KG. Moreover, the filtered KG substantially decreased the time required to optimize and train models, making them more accessible to a broader audience. Training KGE models on large knowledge graphs like Monarch KG is prohibitively resource intensive. For instance, RotatE with the full Monarch KG requires over 30 days to complete hyperparameter optimization on a dedicated NVIDIA A100 80GB GPU. Such computational and time requirements create significant barriers (estimated cost of $3,701.22 on a similar outfitted Google Compute Engine) that restrict KGE model use to institutions with access to shared supercomputer clusters or research computing budgets. Careful filtration of KGs proves to be an effective approach for improving predictive performance while substantially reducing computational time and power requirements, thereby greatly reducing research costs.

Topological imbalance and resulting degree bias have been addressed from multiple angles, including novel computational tools ^36^ and specialized training regimes ^37^. Our results expand this field by highlighting the critical importance of inspecting and curating input data for KGE link prediction. The optimal solution to degree bias will likely require integrating multiple complementary approaches rather than relying on any single methodology ^38^. By combining data curation with state-of-the-art computational and training approaches, we can develop KGE-based link predictors capable of reliable variant prioritization in sparse regions of knowledge graphs, particularly for rare diseases.

## Key Points

- Knowledge graph embedding models exhibit node-degree bias that negatively affects predictions for rare diseases.
- Using a massively aggregate knowledge graph is suboptimal; better performance and runtime can be achieved with selective information filtering.
- The quality of information and edges added to a knowledge graph is more important than the quantity for improving KGE model performance.
- The optimal combination of metanodes should be chosen empirically based on the specific prediction task.

## Study funding

This work was supported by a grant from the Children’s Hospital of Colorado awarded to RL.

## Supporting information

Supplemental Results

## Acknowledgements

The authors thank Clair A Huffine of Insight Illustrations LLC for the creation of the scientific illustrations used in figures 1 and 2.

## Data Availability

This study utilized existing data from the Monarch Knowledge Graph (version 2023-12-16), available at: https://data.monarchinitiative.org/monarch-kg-dev/2023-12-16/index.html. No new data were generated for this research. All code used to perform the analyses and generate results is publicly available on GitHub: https://github.com/MSBradshaw/LinkPrediction.

## Conflict of interest

The authors declare no conflict of interest.

**Michael S. Bradshaw** holds PhD in Computer Science from the University of Colorado Boulder. His main research interests are in graph-based methods for improving our understanding of diseases of genetic origin.

**Anton Avramov** holds PhD in Microbiology, Cell and Molecular biology from Oklahoma State University. He is currently a postdoctoral associate at the Department of Computer Science and Biofrontiers Institute at the University of Colorado Boulder, working for a lab venture startup CodeBreakerTX.

**Alisa Gaskell** holds a PhD in Molecular Microbiology and Biochemistry from the University of East Anglia. She is the Scientific Director of the Precision Diagnostic Lab and Co-chair of Precision Medicine at Children’s Hospital Colorado. Her experience focuses on genomics from both large-scale academic and reference laboratories, with expertise in the development and implementation of next-generation sequencing technologies and the analysis of genomic data. Harmonizing these different approaches, she strives to develop a value-based method to genomics testing that forms the foundational aspect of precision medicine.

**Ryan Layer** holds a PhD in Computer Science from the University of Virginia. He is currently an assistant professor at the Department of Computer Science and Biofrontiers Institute at the University of Colorado Boulder. His research focuses on developing computational methods that can leverage population-scale datasets to better understand how genetic variation affects human health.

## Notes

### Competing Interest Statement

The authors have declared no competing interest.

### Summary of Updates

This version of the manuscript has an improved background section and figures.

