## Supplemental Results for "The effects of biological knowledge graph topology on embedding-based link prediction"

### Methods

| Knowledge Graph | Edges | Nodes |
| --- | --- | --- |
| Monarch KG | 11412471 | 519303 |
| Monarch KG filtered | 1289563 | 43737 |
| HuRI | 97679 | 31579 |
| STRING 100% | 1152555 | 40843 |
| STRING 50% | 608734 | 40094 |
| STRING 25% | 284992 | 37923 |
| CHEBI + Filtered KG | 1301537 | 48660 |
| CHR + Filtered KG | 1289929 | 43940 |
| CL + Filtered KG | 1292776 | 45353 |
| dictyBase + Filtered KG | 1301690 | 48890 |
| FB + Filtered KG | 1593952 | 56464 |
| GO + Filtered KG | 1689454 | 87452 |
| HP + Filtered KG | 1788336 | 60417 |
| HSAPDV + Filtered KG | 1290264 | 43975 |
| MGI + Filtered KG | 1773282 | 64649 |
| NCBIGene + Filtered KG | 2053721 | 86872 |
| NCBITaxon + Filtered KG | 1291965 | 45434 |
| NCIT + Filtered KG | 1289610 | 43758 |
| PATO + Filtered KG | 1290794 | 44368 |
| PomBase + Filtered KG | 1295589 | 46932 |
| Reactome + Filtered KG | 1334419 | 46040 |
| RGD + Filtered KG | 1619008 | 63424 |
| SGD + Filtered KG | 1296195 | 47057 |
| SO + Filtered KG | 1289640 | 43776 |
| UBERON + Filtered KG | 1334609 | 58196 |
| WB + Filtered KG | 1545733 | 57466 |
| Xenbase + Filtered KG | 1440666 | 56984 |
| ZFIN + Filtered KG | 1601905 | 60856 |

**Supplementary Table 1**. **Summary of the 29 filtered knowledge graphs used in this study, showing node and edge counts.** Each filtered KG contains HGNC and MONDO nodes plus one additional metanode type, except for the baseline Monarch KG which includes all original data.

### Hyperparameter optimization

| Parameter | Min | Max | Step |
| --- | --- | --- | --- |
| embedding_dim | 64 | 512 | 16 |
| num_epochs | 100 | 1000 | 100 |
| lr | 0.001 | 0.01 | log |
| num_negs_per_pos | 1 | 100 | 10 |
| loss | NSSA | - | - |
| n_trials | 30 | - | - |
| stopper | early | - | - |
| stopper frequency | 10 | - | - |
| stopper patience | 2 | - | - |
| stopper relative_delta | 0.002 | - | - |

Supplemental Table 2: The parameters and their value ranges searched during hyperparameter optimization. The value log in the step column denotes the value was sampled from the range in the log domain rather than using a discrete step value. A separate hyperparameter optimization pipeline was run for each KG-model pair; the final values for all models used in this study can be found in the supplemental information.

### Sex-specific differentially expressed genes

We use a set of 311 and 352 genes that were differentially expressed between females and males respectively. These gene lists originate from the work of Oliva et al. 2020 (Oliva et al. 2020) in which they identify genes that have sex-based differential expression in the 44 tissues sampled in GTEx. This study reported the top differentially expressed genes in each tissue, we chose the top 100 in each tissue which resulted in sets of 311 and 352 unique genes. Genes were divided into female/male-specific categories based on which sex has greater expression.

### Diseases caused by ancestry-specific variants

We created a set of 1224 diseases caused by an ancestry-specific variation. Broken down by ancestral population there were 195 African, 223 East Asian, 309 Latino, and 497 European. These diseases were identified based on allele frequency (AF) in ancestral population data from gnomAD and variant to disease annotations in OMIM. The criteria for selecting these were:

1. allele with AF != 0
2. allele with pathogenic label
3. allele located in a gene where all alleles that pass criteria 1 and 2 come only from a single ancestry population.

Alleles that passed those criteria were then cross-referenced through OMIM to find diseases with MONDO identifiers caused by that allele.

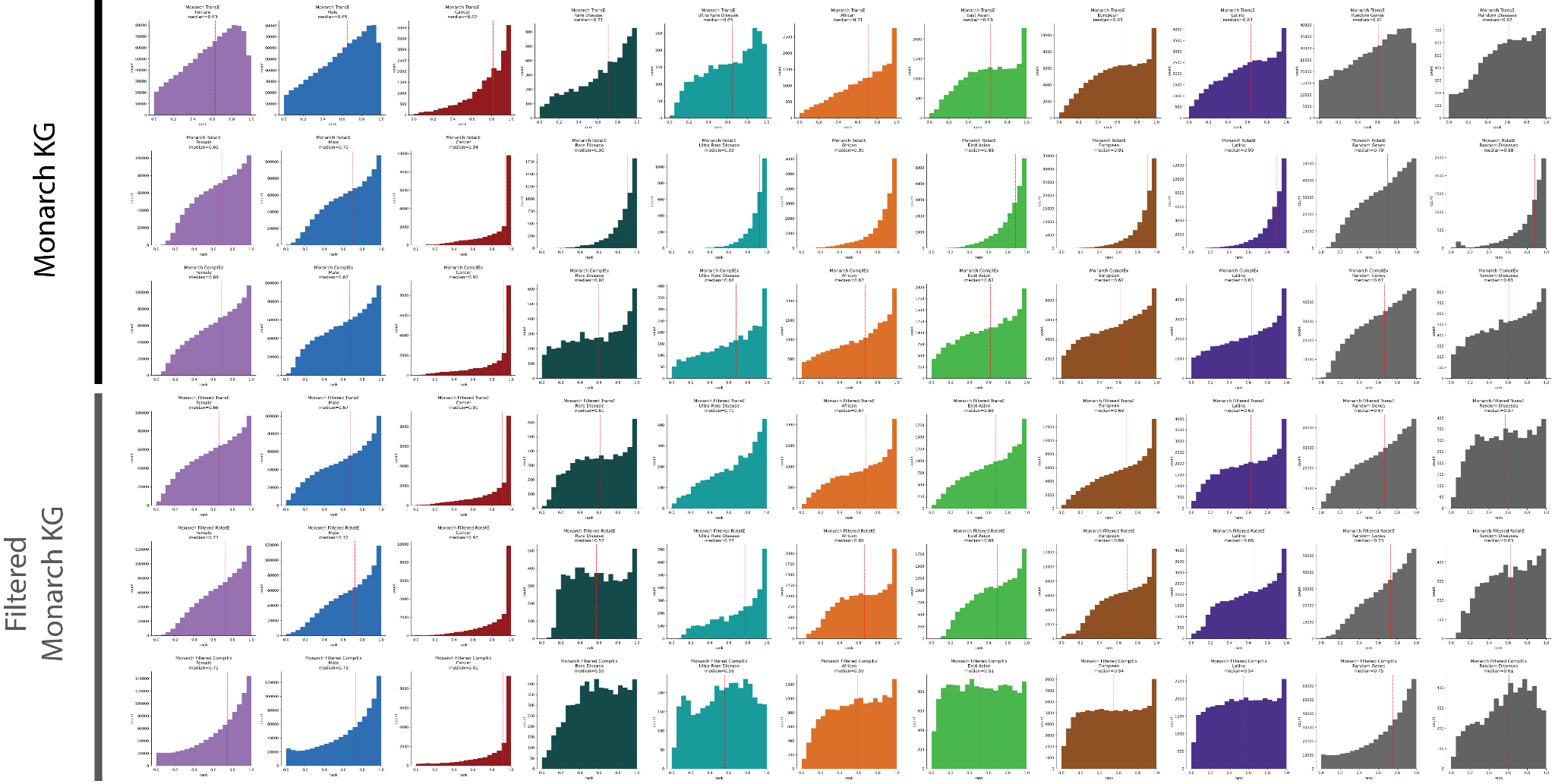

Supplemental Figure 1: Distributions of normalized rank from link prediction on all groups: heritable cancer genes, non-cancer genes, rare diseases, ultra-rare diseases, control diseases, female and male differentially expressed genes, diseases causes by ancestry-specific variants. Results are ordered by the KG used (full Monarch KG or filtered KG) and by KGE model used (TransE, RotatE, ComplEx).

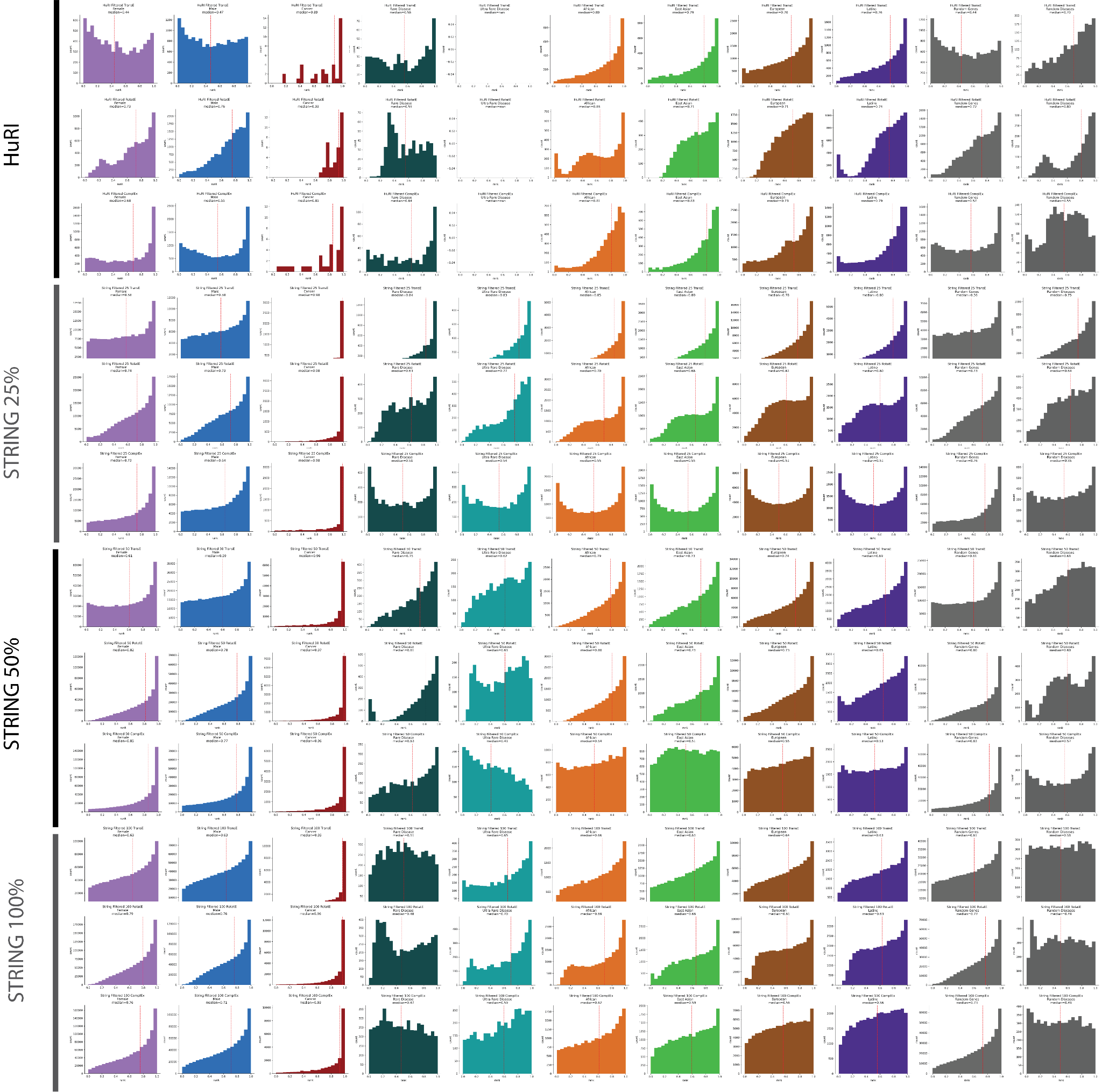

Supplemental Figure 2: Distributions of normalized rank from edge confidence link prediction experiments with all groups: heritable cancer genes, non-cancer genes, rare diseases, ultra-rare diseases, control diseases, female and male differentially expressed genes, diseases causes by ancestry-specific variants. Results are ordered by the KG used (filtered Monarch with PPI replaced by HuRI, STRING top 25%, 50%, or 100%) and by KGE model used (TransE, RotatE, ComplEx).
